# Co-SELECT reveals sequence non-specific contribution of DNA shape to transcription factor binding in vitro

**DOI:** 10.1101/413922

**Authors:** Soumitra Pal, Jan Hoinka, Teresa M. Przytycka

## Abstract

Understanding the principles of DNA binding by transcription factors (TFs) is of primary importance for studying gene regulation. Recently, several lines of evidence suggested that both DNA sequence and shape contribute to TF binding. However, the question if in the absence of any sequence similarity to the binding motif, DNA shape can still increase probability of binding was yet to be addressed.

To address this challenge, we developed Co-SELECT, a computational approach to analyze the results of *in vitro* HT-SELEX experiments for TF-DNA binding. Specifically, the presence of motif-free sequences in late HT-SELEX rounds and their enrichment in weak binders allowed us to detect evidence for the role of DNA shape features in TF binding.

Our approach revealed that, even in the absence of the sequence motif, TFs have propensity to weakly bind to DNA molecules enriched in specific shape features. Surprisingly, we also found that some properties of DNA shape contribute to promiscuous binding of all tested TF families. Strikingly, such promiscuously bound shapes correspond to the most frequent shape formed by the DNA. We propose that this promiscuous binding facilitates diffusing of TFs along the DNA molecule before it is locked in its binding site.

## 1 Introduction

Characterizing the DNA binding specificities of transcription factors (TFs) is a fundamental step in studying gene regulation. It is well established that transcription factors typically bind to specific sequence motifs [1, 2]. An increasing amount of studies additionally suggest that the nucleotides adjacent to the binding motifs also contribute to the TF-target interactions [3, 4]. Since these residues do not make contact with the TF binding site, it is assumed that their contribution to the binding specificity is indirect. Recently, it has been proposed that this contribution is achieved by influencing the structural properties of the DNA molecule. These shape related DNA characteristics include the minor grove width, roll, propeller twist, and helical twist [5, 6, 7, 8].

Several lines of evidence indicate that DNA structure is important for TF binding. First, there are examples of co-crystal structures that show complementarity of 3D structures of TF and DNA [6, 9, 10, 11]. However, the number of such co-crystal structures is relatively limited and conformational changes upon binding might also contribute to this complementarity. Next, DNA shape has been found to possess significant effects on the electrostatic potentials near the surface of the molecules, particularly in the grooves [5]. Indeed, recent computational analyses revealed correlation between minor-groove width and electrostatic potential, although identity of nucleotides also plays some role in the manifestation of the potential [12]. As further support, using hydroxyl radical cleavage as a measure of local DNA structure, a computational study [13] on the ENCODE regions of human genome found common DNA structural motifs apparently without any sequence consensus in a large collection of DNase I hypersensitive sites that are closely associated with the regulatory elements. In addition, a recent study that considered a family of TFs and analyzed the DNA shape of thousands of their binding sites, demonstrated covariation between the protein sequence and both the sequence and shape of their DNA targets [14]. Finally, several computational analyses used machine learning approaches to show that using shape features improves prediction of TF binding both, *in vitro* [15, 16, 17, 18] and *in vivo* [11, 16, 17]. In addition, [19] developed a method to identify shape motifs from *in vivo* biding data.

Despite significant effort, evidence for a role of shape for TF binding is mostly indirect. One of the principal challenges in measuring this effect directly is related to the fact that DNA shape is primarily determined by its underlaying sequence. This interdependency in turn substantially aggravates the task of disambiguating the individual contribution of sequence and structure. Indeed, computational inference of DNA shape features is based on sequence information alone and thus computationally inferred shape features could be seen as functions summarizing sequence information of the neighboring residuals [20, 21]. Consistent with this view, machine learning models for TF-DNA binding have been reported to show a similar improvement when features based on two or three consecutive nucleotides in lieu of shape features were added [15, 18, 22, 23]. In particular, a recent study showed that sequence-to-shape conversion can be estimated nearly perfectly based on mononucleotide and dinucleotide features [22, 23].

These results raise the question whether shape can be deconvoluted from sequence so that the contribution of shape alone could be tested. In light of the above findings, an independent shape contribution could only be assessed by testing for binding to a DNA molecule that is completely free of any similarity to the sequence binding motif but does contain the shape corresponding to the motif sequence. To obtain such evidence we reexamined the data obtained through a collection of HT-SELEX experiments on TF binding [18] using a novel approach that we developed for this purpose and refer to as Co-SELECT.

HT-SELEX has been extensively used to study *in vitro* RNA and DNA binding. This experimental technique leverages the paradigm of *in vitro* selection by repetitively enriching a pool of initially random RNA/DNA sequences with those that bind a target of interest [24, 25, 26]. At the end of the selection process, the sequences that are bound to the target are expected to be present in multiple copies, while remaining sequences consist of weak binders and/or noise and are typically excluded from further analysis or broadly considered as “background”.

In the context of our studies, HT-SELEX data provides several unique advantages. First, selection is performed *in vitro* and thus binding events are not influenced by other complex DNA transactions (such as transcription, replication, etc) that take place in the cell. For example, gene transcription is typically accompanied by negative supercoiling of the DNA molecule upstream of the transcription start site which, in turn, might critically influence DNA shape [13, 27, 28]. Next, while the initial pool of DNA fragments is assumed to contain random DNA sequences, intermediate pools are increasingly non-random as the strongest binders are amplified and non-binders eliminated. Weaker binders would eventually be out-competed given a sufficient number of selection rounds. These species however can survive the section process and are expected to be enriched among the background sequences since the number of selection rounds of SELEX targeting TFs is typically low.

To leverage the properties of *in vitro* selection for the purpose of investigating the possibility of selection for shape features, a novel approach capable of capturing such features is needed. Here, we introduce such an approach that utilizes the concept of a shapemer. A shapemer is similar to that of a *k*-mer but rather than being a sequence of nucleotides, a shapemer represents a sequence of shape features at nucleotide resolution. Based on this concept, we developed Co-SELECT - a method for detecting shapemers enriched in both: oligos that contain binding motifs and the background oligos that survived until the final round. Note that unlike previous studies which aimed at addressing the question of whether computationally derived shape features of the binding motif provide additional information relative to the sequence alone, we ask if in the absence of any sequence similarity to the binding motif, can shape alone provide any binding advantage? Our results demonstrate that this is indeed the case. Interestingly, we also found promiscuous shapemers that are favored in non-specific TF binding.

## 2 Materials and Methods

### 2.1 Data

In this study we utilize the HT-SELEX datasets from [18] which is derived from the experiments in Jolma et al. [26] and complemented with additional sequencing data. To the best of our knowledge, this dataset is currently the most extensive mammalian TF-DNA binding dataset derived using HT-SELEX [18]. Here we are focusing on the following three families of transcription factors: 1) basic helix-loop-helix (bHLH), 2) E26 transformation-specific (ETS) and 3) homeodomain. Our choice of the three families is motivated by the fact that these families are assumed to have a uniquely defined, very highly conserved *core-motif*. Our analysis depends on such well defined core-motifs for a clear classification of oligos into three categories depending on the presence and completeness of the core-motif as described below. The default core-motifs for the three families are CACGTG, CGGAA and `TAAT`, respectively [18], however for each TF we also tested whether the selection results are consistent with the default core-motif, and if not, an alternative motif of the same length derived in [26] from the HT-SELEX results was also considered (Table S1 in Supplementary Section S1).

Finally, Jolma et al. observed that some of their experiments show evidence of cross contamination artifacts or indication of an unsuccessful selection [29]. Consequently, we refined our dataset by removing experiments with the above described signatures from further analysis based on the general strategy outlined in that paper. Since our method relies on data from experiments that generated a strong core-motif, we further discarded those selections which yielded ambiguous binding motifs (Supplementary Section S1). After this filtering, out of the initial set of 212 datasets, 131 high quality datasets remained which are comprised of 14 experiments in the bHLH family, 18 experiments in the ETS family, and 99 experiments in the homeodomain family (Table S1 in Supplementary Section S1).

### 2.2 HT-SELEX procedure and classification of oligos with respect to motif presence

Our approach to analyzing the contribution of DNA shape on TF binding leverages the HT-SELEX (High-Throughput Systematic Evolution of Ligands by EXponential enrichment) protocol and resulting sequencing data. Since the basic properties of the HT-SELEX protocol are fundamental to our method we start with a brief summary of the relevant features of HT-SELEX. In a nutshell, HT-SELEX is an iterative procedure that starts with an initial pool of random DNA sequences (also refereed to as *oligonucleotides* or *oligos*) which are typically 20-40 nucleotides in length.

Each iteration of HT-SELEX can be viewed as a competition among the oligos for binding to the TF. Oligos that do not bind at all or bind weakly are washed out from the pool while the remaining species are amplified. A sample of the amplified pool is sequenced to allow for computational analysis while another sample is used as the input for the subsequent selection round. In this way, the proportion of high-affinity oligos in the pool increases at each iteration while non-binders and, subsequently, weaker binders are gradually eliminated/out-competed. All the oligos in the final round are generally called *aptamers* as they are expected to be the true binders for the target TF.

We say that an oligo is *motif-containing* if it contains the exact core-motif sequence, 2) *motif-free* if it does not share more than 2 nucleotides with the core-motif at any position, and 3) *partial-motif-containing* otherwise. The threshold 2 was selected as the largest value *d* such that a random oligo of length *n* has, with probability ≥ 0.9, at least *d* nucleotides common with the core-motif. For the smallest length of an oligo in our datasets (*n* = 20) and the shortest core-motif (*k* = 4) the probability of overlap on *d* = 1, 2, 3 positions are estimated as 0.99, 0.98, 0.56 respectively (Supplementary Section S2). Thus, the threshold *d* = 2 (but not *d* = 3) for the maximum number of bases matching with the core-motif ensures that a motif-free oligo can be safely assumed to carry no sequence information of the core-motif.

Importantly, analysis of the sequencing data from consecutive iterations can provide key information about the dynamics of this competition as illustrated in Figure 1. Using simulated data generated with AptaSim [30], Figure 1A shows how the proportions of five groups of oligos with varying affinity to the target change throughout the selection process. Initially, the proportion of medium and high affinity binders increases at the expense of weak binders and non-binders. However, in later rounds the proportion of medium binders starts to decrease indicating that they too are now out-competed by stronger binders. At this point the proportion of non-binders is drastically reduced and quickly becomes negligible.

**Figure 1:**
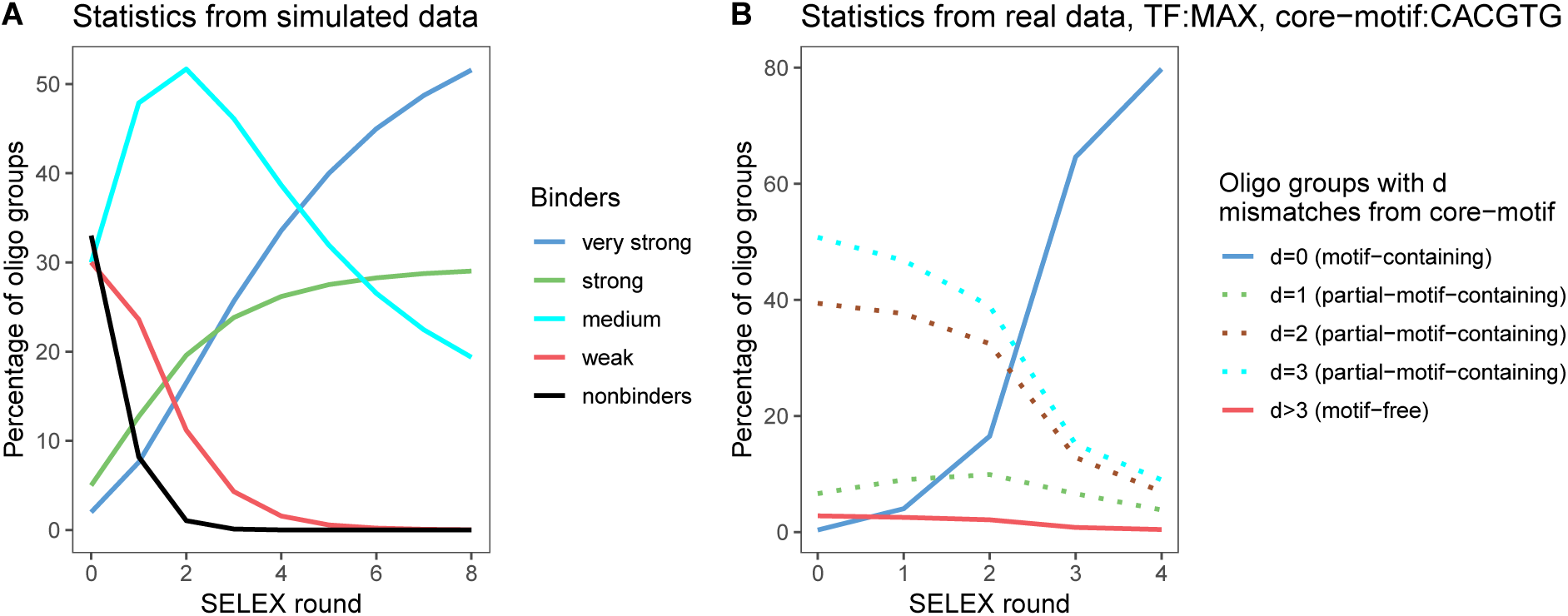
Competition between binders using simulated and real HT-SELEX data. A. Simulation of 8 rounds of HT-SELEX using AptaSim[30] where the initial pool contained 10 million unique oligoes divided into 5 groups where binding probabilities withing each group are sampled from a differently parametrized normal distribution: very strong binders (mean 0.90 and SD 0.1), strong binders (mean 0.50, SD 0.30), medium binders (mean 0.25 SD 0.25), weak binders (mean 0.1, SD 0.1) and non-binders (mean 0.025, SD 0.025). The curves show the distributions of the four groups of oligos across the HT-SELEX rounds. B. The selection dynamics for the transcription factor MAX. The oligos are categorized into three groups depending on the best match to the core-motif CACGTG. The motif-free group is expected to be a mixture of nonbinders and weak background binders.

These expected trends are indeed observed in experimental data as exemplified by the selection results performed against the transcription factor myc-associated factor X (MAX) in Figure 1B. The figure demonstrates a rapid increase of the population of oligos that contain the complete core-motif, consistent with the expectation that this group contains the strongest binders. The populations of oligos that contain partial-motifs with one mismatch from the core-motif initially increases as expected from medium strength binders while population of oligos containing partial motifs with 2 or 3 mismatches and the motif-free oligos are out-competed. Given four rounds of selection, even the oligos in the motif-free group that survive this competition are expected to be enriched in (presumably weak) binders.

### 2.3 Discretized shape strings and shapemers

We use the term *shapemers* to describe constant length sequences of discretized shape values for a specific DNA feature such minor groove width (MGW), propeller twist (ProT), helical twist (HelT), and Roll. We utilized DNAShape [21], which given a DNA sequence, uses a sliding-window approach to estimate DNA shape at each position. For the purpose of this study, we discretized the shape values using cutoffs based on the frequency distribution of the shape values in the oligos from the initial pools (Supplementary Section S3). Two different sets of discretization thresholds were examined to confirm that the conclusions drawn do not depend on these threshold values. We used four discretization levels for MGW, ProT and Roll denoted as S (Small), M (Medium), H (High), X (eXtra high) and three levels for HelT (S,M,H) (Supplementary Section S3). In this study we used shapemers of length 6. This length was selected as it corresponds to the length of the longest core-motif considered in this study.

We analyze the shapemers from the two groups of oligos, motif-containing and motif-free, separately. While the shapemers from the motif-containing oligos carry the sequence information of the core-motif, the shapemers from the motif-free oligos are devoid of any sequence information of the core-motif. Indeed, we confirmed that shapemers enriched within the subpopulation of motif-free aptamers do not share sequence similarity with the core-motif (Supplementary Section S4).

### 2.4 Computing enrichment of shapemers

In this paper, we use fold enrichment to denote the ratio between the frequency of a particular feature in the final SELEX pool to its frequency in the initial pool. Here we focus on the enrichment of shapemers and use a Markov model to estimate the expected presence of shapemers in the initial pool. To analyze the enrichment of shapemers in motif-specific binding we consider only the shapemers that are contained in the interval consisting of the core-motif and a flanking nucleotide on either side (Figure 2). For motif-free binding we do not have any prior assumption which shapemer might be involved in binding and thus we consider all shapemers contained in the apatmer. We denote the shapemers thus identified from the motif-containing and motif-free oligos as *core shapemers* and *motif-free shapemers* respectively.

**Figure 2:**
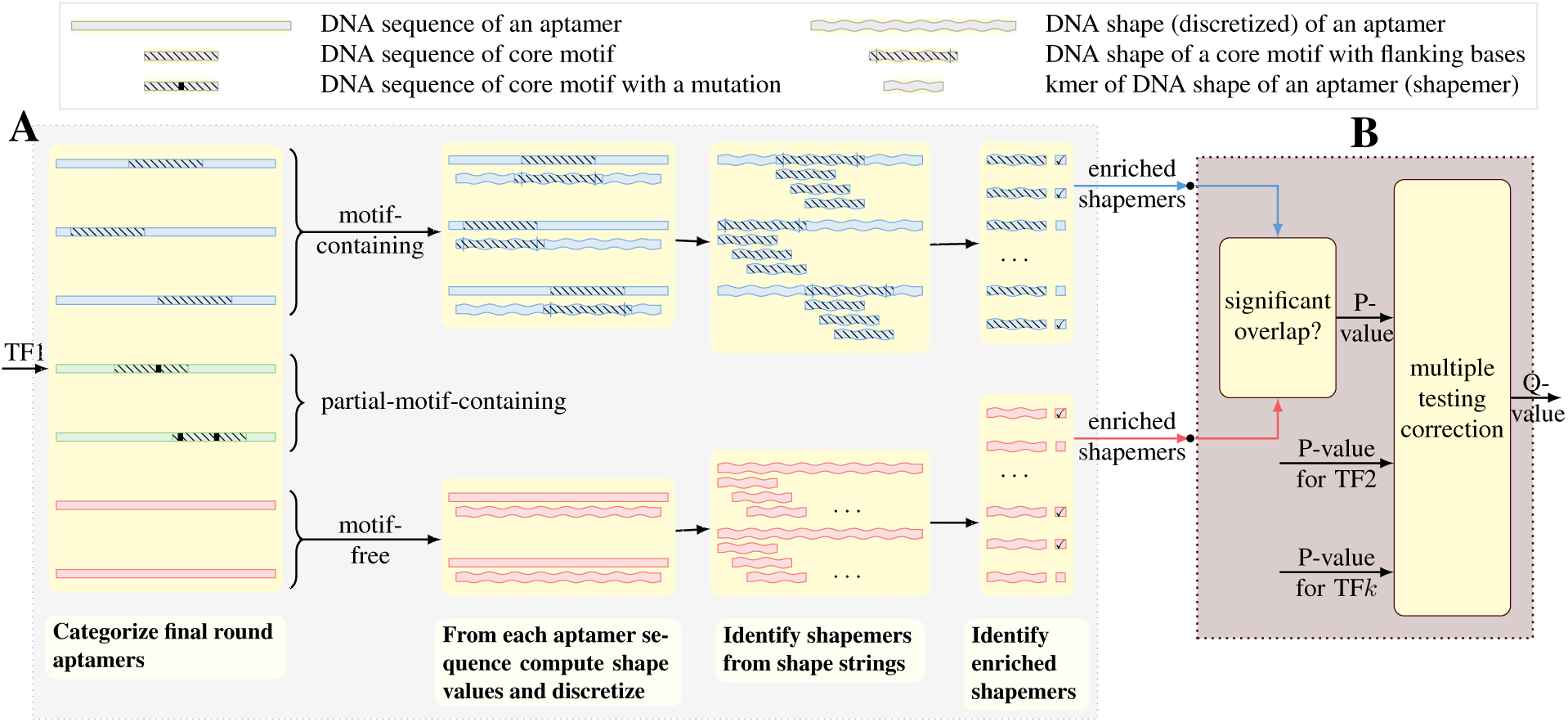
Overview of the Co-SELECT method. A. The aptamers present in the last round are categorized into three types: 1) motif-containing, 2) partial-motif-containing and 3) motif-free (see Material and Methods). Next, the shapemers that potentially contribute to binding in the motif-containing group (core shapemers) and in the motif-free aptamers are identified. B. Testing the significance of the overlap of enriched core and motif-free shapemers.

Let *j* ∈ {*m n*} index the two sets of shapemers of type core and motif-free respectively. We compute *R*_*j*_ (*x*), the enrichment of shapemer *x* in set *j* as

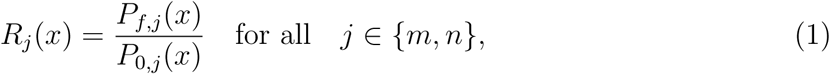

where *P*_*i,j*_ denotes the probability of a shapemer *x* appearing in set *j* and round *i*. We compute the probabilities for the final round *f* as

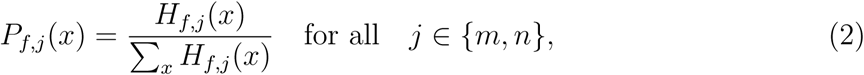

where *H*_*f,j*_ (*x*) denotes the number of times shapemer *x* appears in set *j* and round *f*.

We estimate the probabilities *P*_0,*j*_(*x*) for the initial pool using a Markov model, the details of which are given in Supplementary Section S6. We discard the shapemers *x* with *R*_*j*_(*x*) *< ρ* where *ρ* denotes the *enrichment threshold*. We experimented with four values of *ρ* ={1.05, 1.10, 1.20, 1.50} and report the results for *ρ* = 1.2 since it resulted in a lower false discovery rate relative to the smaller thresholds while the enrichment ≥ 1.5 was very rare. In addition, to reduce noise, we only consider shapemers that are present in at least 1% of aptamers (Supplementary Section S5).

### 2.5 Statistical analysis

We assess the statistical significance of the overlap of enriched motif specific and motif free shapemers in the last round of selection using the one-sided Fisher’s exact test and p-value cut-off 0.05. To correct for multiple testing we used the q-value method [31]. To compare with the control experiment (see Results), we use *ECR*_*p*_ defined as the fraction of the statistically significant experimental results with p-value cut-off *p* to the fraction of the control results with the same p-value cut-off. To avoid division by zero, we add 10^-5^ to the fraction for control groups.

## 3 Results

### 3.1 The Co-SELECT method for detecting shapemers present in motif-containing and motif-free aptamers

We reasoned that if the DNA shape contributes to TF binding in a sequence independent way, then after several selection rounds we should observe enrichment of the same shape features in both motif-specific and non-motif-specific binding. With this in mind, we developed our Co-SELECT method to test for such simultaneous selection for the same shape features in two extreme subpopulations of oligos from the same HT-SELEX experiment. The first subpopulation corresponds to the set of aptamers that contain the core binding motif without any mismatches and the second subpopulation encompasses aptamers that have no similarity to the core-motif. To describe shape features we use *shapemers* - constant length sequences of discretized values of minor groove width (MGW), propeller twist (ProT), helical twist (HelT), or Roll. If both groups show significant enrichment in the same shapemers, such co-enrichment cannot be attributed to sequence similarity but rather to a selection for these shapemers in both populations.

The graphical overview of the Co-SELECT method is shown in Figure 2A while the technical details related to selecting the two extreme population, computing the enrichment of shapemers in both of them are described in the Materials and Methods section. In brief, first the aptamers present in the last round are categorized into three groups: 1) with complete motif (motif-containing) 2) with partial motif, and 3) motif-free. Next Co-SELECT identifies, the shapemers that potentially contribute to binding in the (complete) motif-containing and the motif-free groups. Finally, the enrichment of shapemers in both groups is computed and tested using Fisher test for statistically significant overlap and subsequently corrected for multiple testing (Materials and Methods).

### 3.2 Co-SELECT identifies DNA shapemers that contribute to the strength of motif-specific and non-motif specific binding

We applied Co-SELECT to the HT-SELEX data that passed the quality control described in the Methods section focusing on three large families of transcription factors; ETS, bHLH and homeodomain. These three families are structurally different and contain a strongly conserved core-motif whose lengths differs between the families. Overall, we observed statistically significant selection for shape for some members in each family. After multiple testing correction however, the results for homeodomain were no longer found to be statistically significant (Table 1). In addition, our results indicate that the dominant shape features that simultaneously contribute to both motif-specific and non-motif-specific binding are related to MGW, HelT and Roll while ProT shapemers were found to be statistically significant only for the bHLH family.

**Table 1:** Summary of Co-SELECT results. For each combination of shape feature and family we compute the Q-values at p-value of 0.05 or in short *qvalue*_0.05_. The results with *q*-value less than 0.15 are in bold.

| $q_{0.05}$ | **bHLH** | **ETS** | **homeodomain** |
| --- | --- | --- | --- |
| MGW | **0.025** | **0.110** | 0.230 |
| HelT | **0.014** | **0.034** | 0.421 |
| ProT | **0.048** | 1.000 | 0.283 |
| Roll | **0.114** | **0.045** | 0.910 |

As mentioned above, Co-SELECT uses Fisher’s exact test to assess the significance of the overlap between enriched core shapemers and motif-free shapemers from the same experiment. It is theoretically possible that some shapemers are generally enriched in motif-free groups independently of the targeted transcription factor. If such shapemers exists they could inflate statistical significance of the overlap. To exclude this possibility we performed an additional control by applying Co-SELECT after swapping the subpopulations of motif-containing and motif-free aptamers between pairs of transcription factors (Figure 3). Specifically, using *TF*_*m*_ and *TF*_*nm*_ to denote respectively the subpopulation of motif-containing and motif-free aptamers from the last selection round for a transcription factor TF, we tested for co-selection of shapemers in 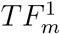 and 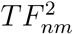 where *TF* ^1^ and *TF* ^2^ are transcription factors from two different transcription factor families. We used all 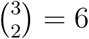 possible pairs of families and all pairs of TF within the respective pair of families. To compare the results of Co-SELECT from the original selection experiments to the so designed controls, we compared the p-value distributions for each TF family and shape (Figure 4) using the experiment to control ratio at the p-value *ECR*_*p*_ (see Materials and Methods). The results were consistent with the results summarized in (Table 1) and provide evidence that the co-selection for shape in the ETS and bHLH families is not an artifact of the experimental procedure.

**Figure 4:**
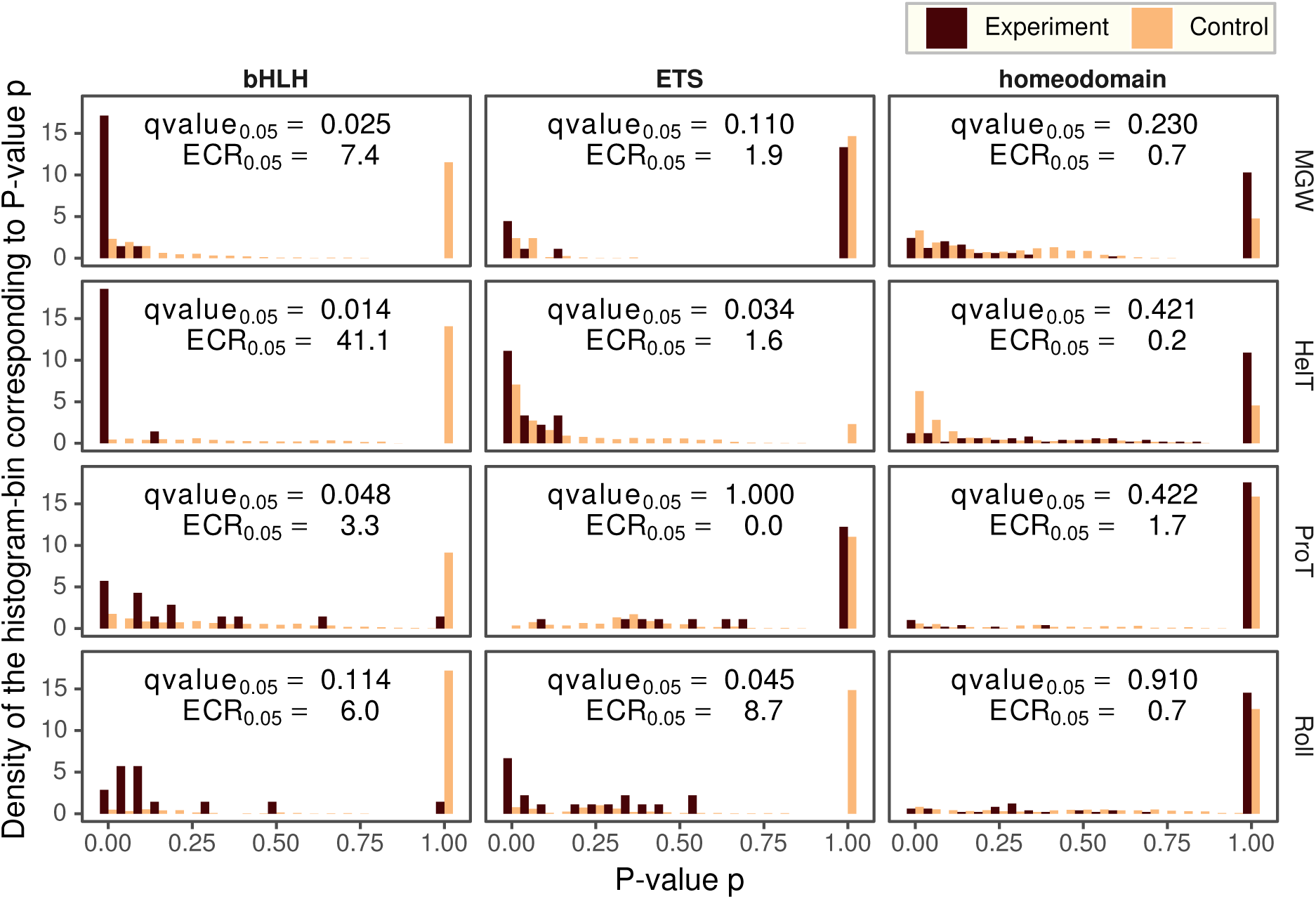
Comparison of p-value histograms for the original and the control experiments.

**Figure 3:**
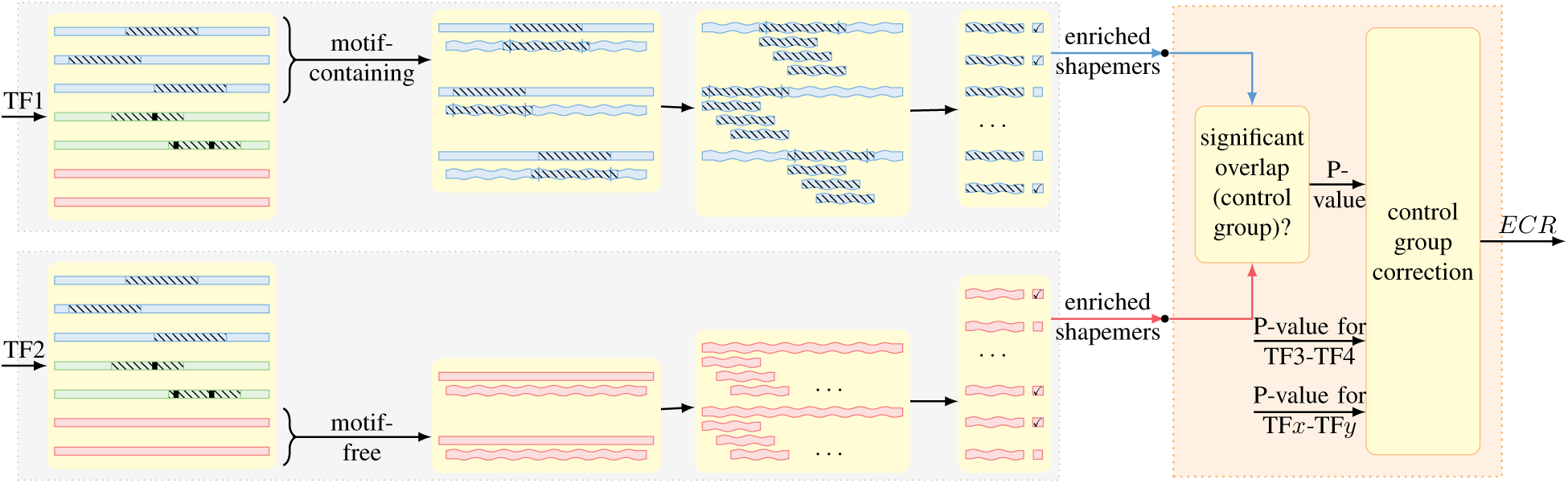
The design of control Co-SELEX experiments. Given two transcription factors (TF1, TF2) from two different families we test for the significance of the overlap of enriched core shapemers from the selection experiment for TF1 and enriched motif-free shapemers from the selection experiment for TF2.

### 3.3 Promiscuity of motif-free shapemers

Next, we asked whether there exist shapemers that are generally enriched in motif-free groups independently of the targeted TF. If such promiscuous shapemers exist, they could provide information about a possible mechanism for non-specific background binding to DNA. To measure promiscuity of a shapemer we compute, for each family, the fraction of TFs where the shapemer is enriched in the motif-free group. The promiscuity of the shapemer is then defined to be the minimum of the so computed fractions over all families. Figure 5 elucidates the shapemers sorted by their promiscuity measures with the identities of the most promiscuous shapemers shown in the insets.

**Figure 5:**
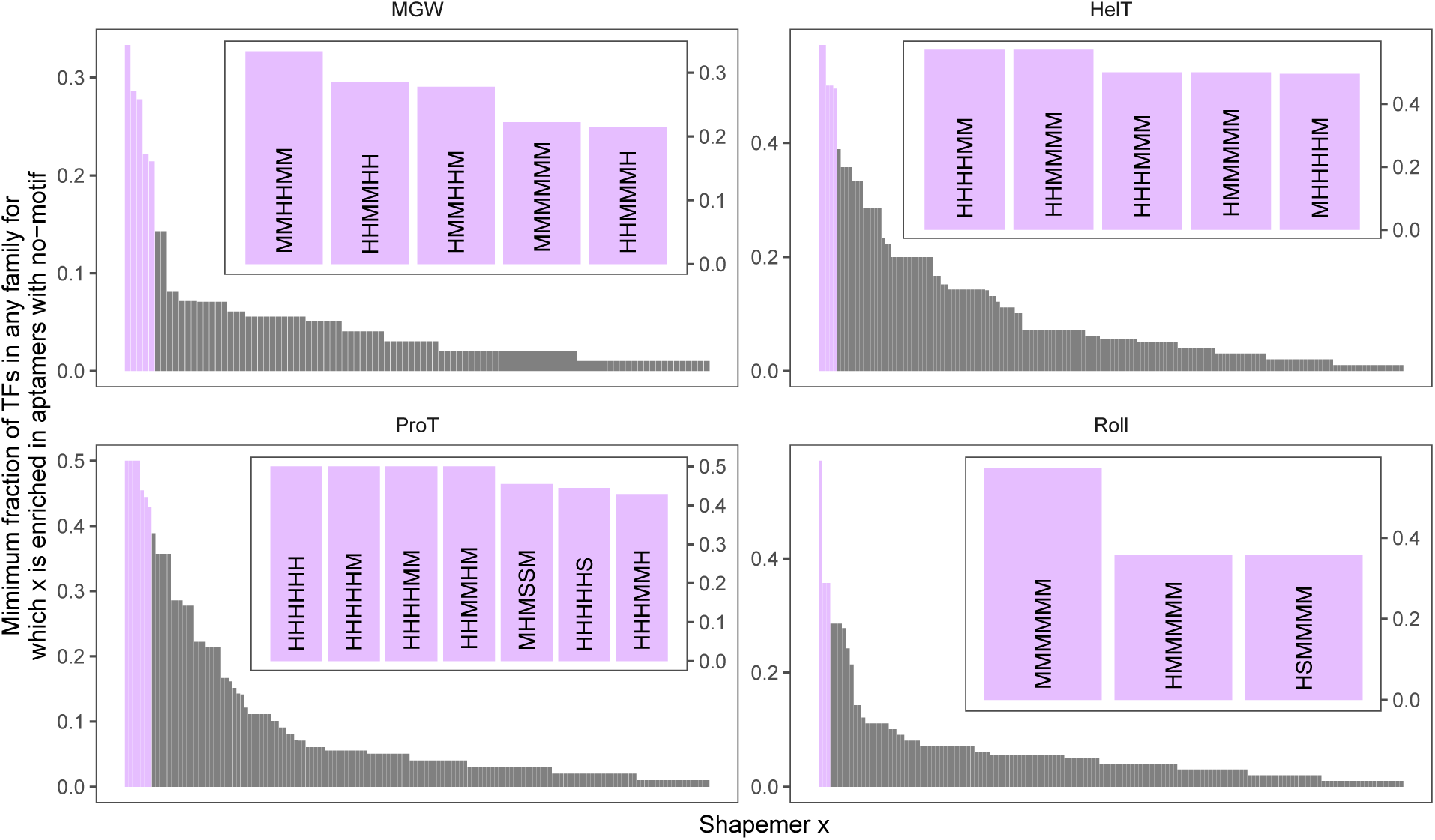
Promiscuity of shapemers. The shapemers are sorted in the decreasing order of their promiscuity. In the inset we show most promiscuous shapemers.

To better understand the source of the observed promiscuity, we considered the fact that the mapping between DNA sequence *k*-mers and shapemers is many-to-one and that the number of sequence *k*-mers mappings to a given shapemer vary between shapemers. To test if the promiscuity is not simply the result of a large number of sequence *k*-mers mapping to a given shapemer, we performed a permutation test. Specifically, for each shapemer we first computed the number of sequence *k*-mers that map to the shapemer and then randomly selected the same number of sequence *k*-mers from the sequence *k*-mers that do map to the shapemer. To compute a p-value we repeated this step 1000 times. We found that the existence of the promiscuous shapemers as shown in the inset of Figure 5 cannot be explained by a large number of sequence *k*-mers mapping to a given shapemer, and are statistically significant with p-value < 10^-3^.

## 4 Discussion

Several previous studies provided evidence that including shape as one of the features in programs that predict biding affinity given DNA sequence, improves the results of such prediction [11, 15, 16, 17, 19]. These results suggest that DNA shape is important for TF binding. However, the question if in the absence of any sequence similarity to the binding motif, can DNA shape still increase probability of binding was yet to be addressed.

The Co-SELECT approach is carefully designed to examine this question. We focused on TFs with very strong and well defined core-motifs allowing us to confidently identify a motif-free population of aptamers. Using two different tests we demonstrated a robust motif-independent contribution of shape-dependent binding. It is possible that for transcription factors with less conserved binding motif, shape might have an even higher contribution to binding. For example, conservation for shape might dictate substitution rules in sequence binding motives. It is also important to point out that our stringent criterion for calling oligos to be motif-free drastically constrains the set of aptamers that are in this category.

While allowing for rigorous analysis, it could reduce its statistical power. This opens the possibility that the impact of shape for motif-free biding not only exists but is even more prominent than estimated in this work.

Interestingly, we identified shapemers that might facilitate weak, non-specific binding. This allows for formulating hypotheses about their roles in TF binding. Focusing on MGW we note that the promiscuously preferred shapes are biased towards M (medium) and H (high) categories. These shape values occur frequently in the normal configuration of DNA as seen in the frequency distributions of shape values in Supplementary Figure S2. This suggests that TF-DNA binding might have evolved to have an affinity to non-specifically bind DNA in its most abundant (favored by many sequences) shape. Such non-specific affinity to the generic DNA shape might facilitate diffusion/sliding of a TF along the DNA molecule as proposed to model for TF binding [32, 33, 34, 35, 36, 37].

## Availability

The raw sequencing data from the HT-SELEX experiments are available in the ENA (http://www.ebi.ac.uk/ena) under study identifier PRJEB14744. The source code for Co-SELECT is available on the public Github repository https://github.com/ncbi/Co-SELECT.

## Acknowledgments

This research was supported by the Intramural Research Program of the National Library of Medicine.

## Supplemental Materials

### S1 Filtering experiments

We used two simple tests to determine if the experiment was successful. The first test, denoted here as *competition-test*, requires that at least 30 percent of the aptamers in the final round must contain the core-motif exactly. The second test that we call *sample-enrichment-test* requires that at least 50 percent of aptamers, that were among the top 100 enriched aptamers in round 3 of SELEX, must retain the enrichment level in round 4 as well. The enrichment-test is a simplified version of a test used in [29] to detect if there was contamination from nearby lanes during the sequencing step of the experiment. The competition-test ensures that the pool of the final SELEX round had enough strong binders, as we saw in the simulation in Figure 1A. The competition test also helps us to determine the core-motif that we use in our analysis. We choose the candidate that results in the maximum percentage of aptamers containing it in the final SELEX round. The results of the two tests on the experiment for ETS family transcription factor ELF5 is shown in Supplementary Figure S1.

**Figure S1:**
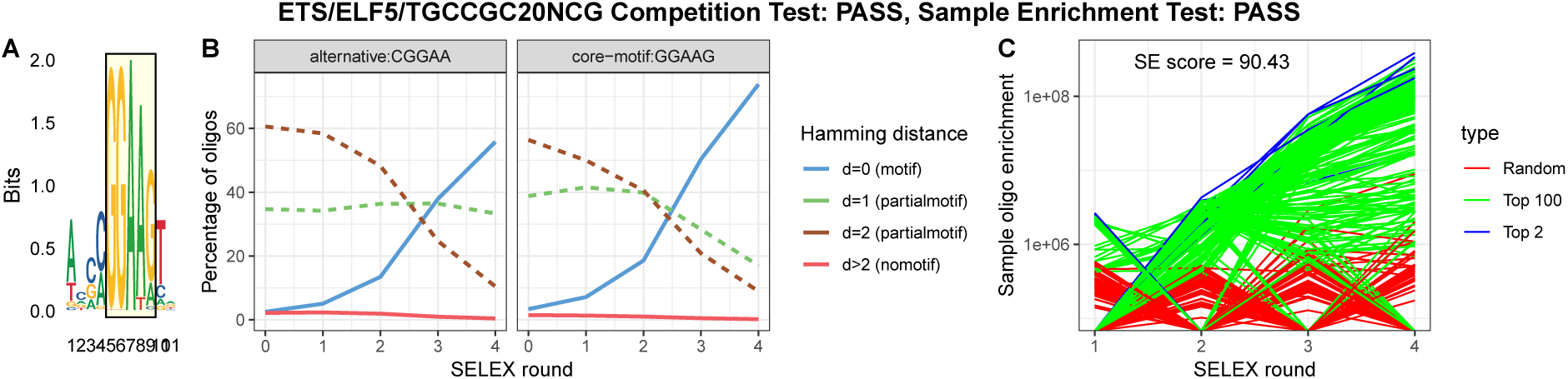
Results of the competition-test and sample-enrichment-test for the experiment for ELF5 from ETS family. A. Most conserved 5 consecutive bases of sequence logo from [26] shows GGAAG. B. For both choices of core-motifs CGGAA (our initial choice for ETS family) and GGAAG, percentage of motif-containing aptamers in the final round is more than 30 and hence the experiment passes competition-test. We use GGAAGG as core-motif in subsequent analysis as it gives larger percentage of motif-containing aptamers. C. SE score = 90.43, i.e., among the 100 most enriched aptamers from each round, only 90.43 percent maintained the enrichment level from round 3 to round 4. Since this score is more than 50, the experiment passes sample-enrichment-test too.

**Table S1:**
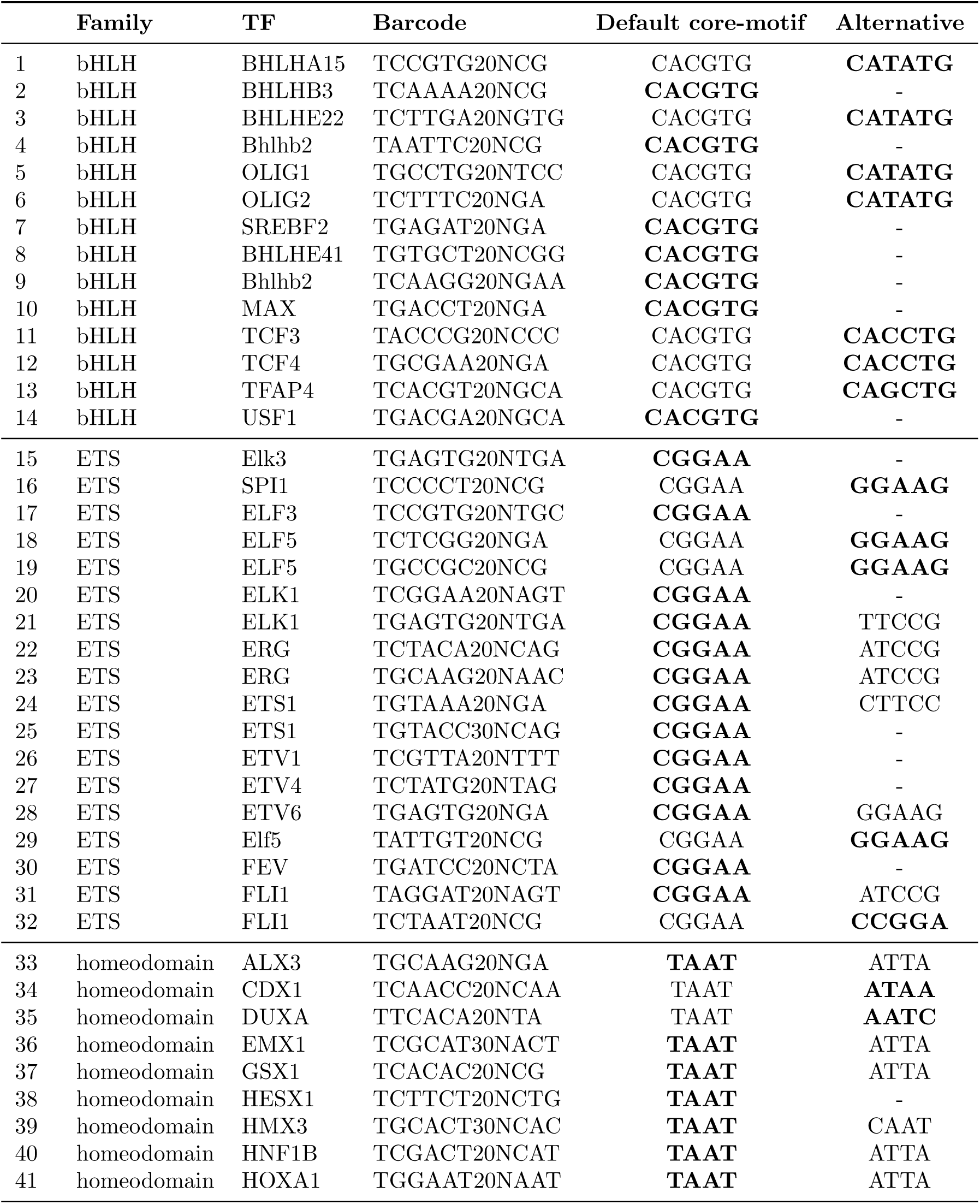
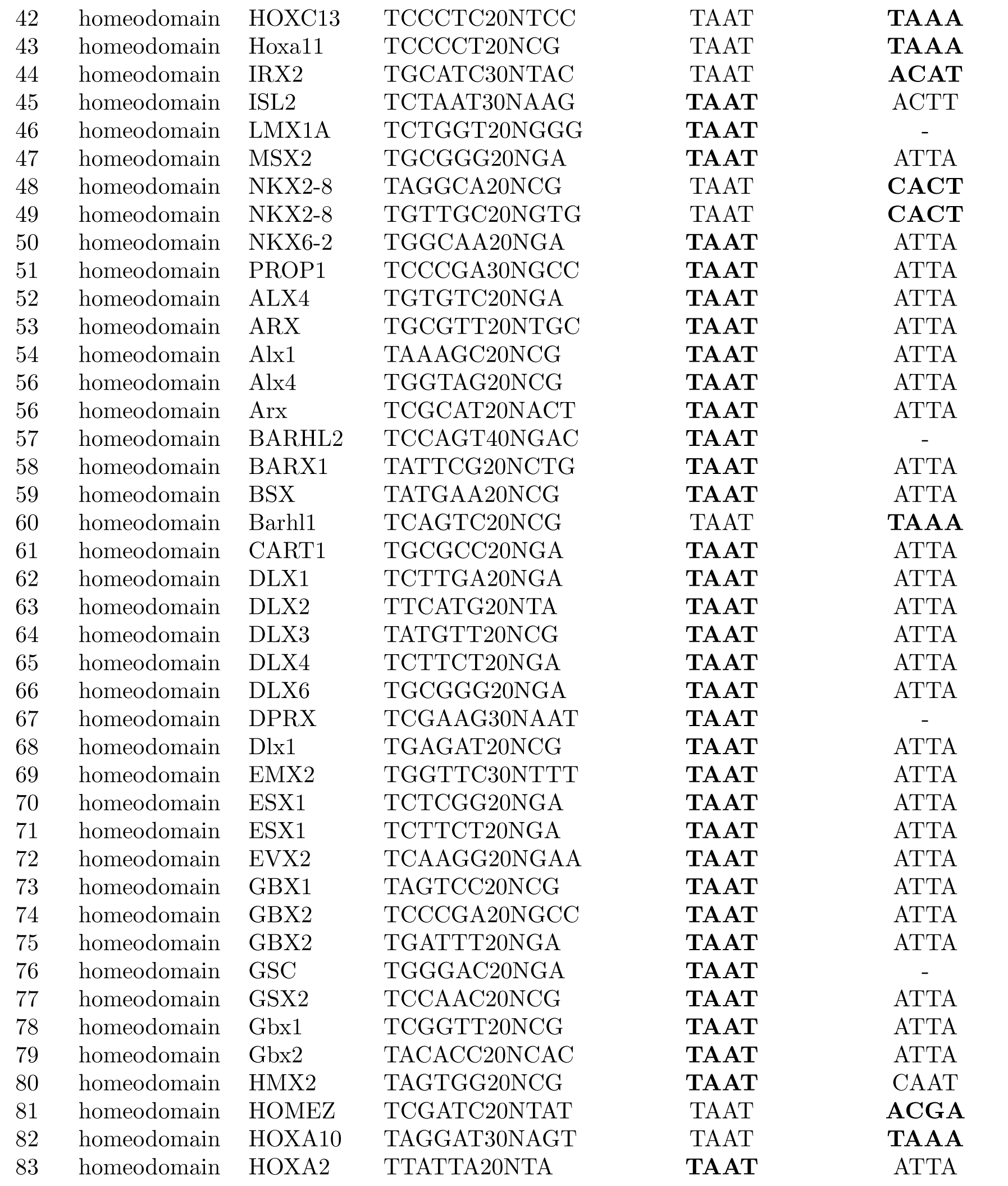
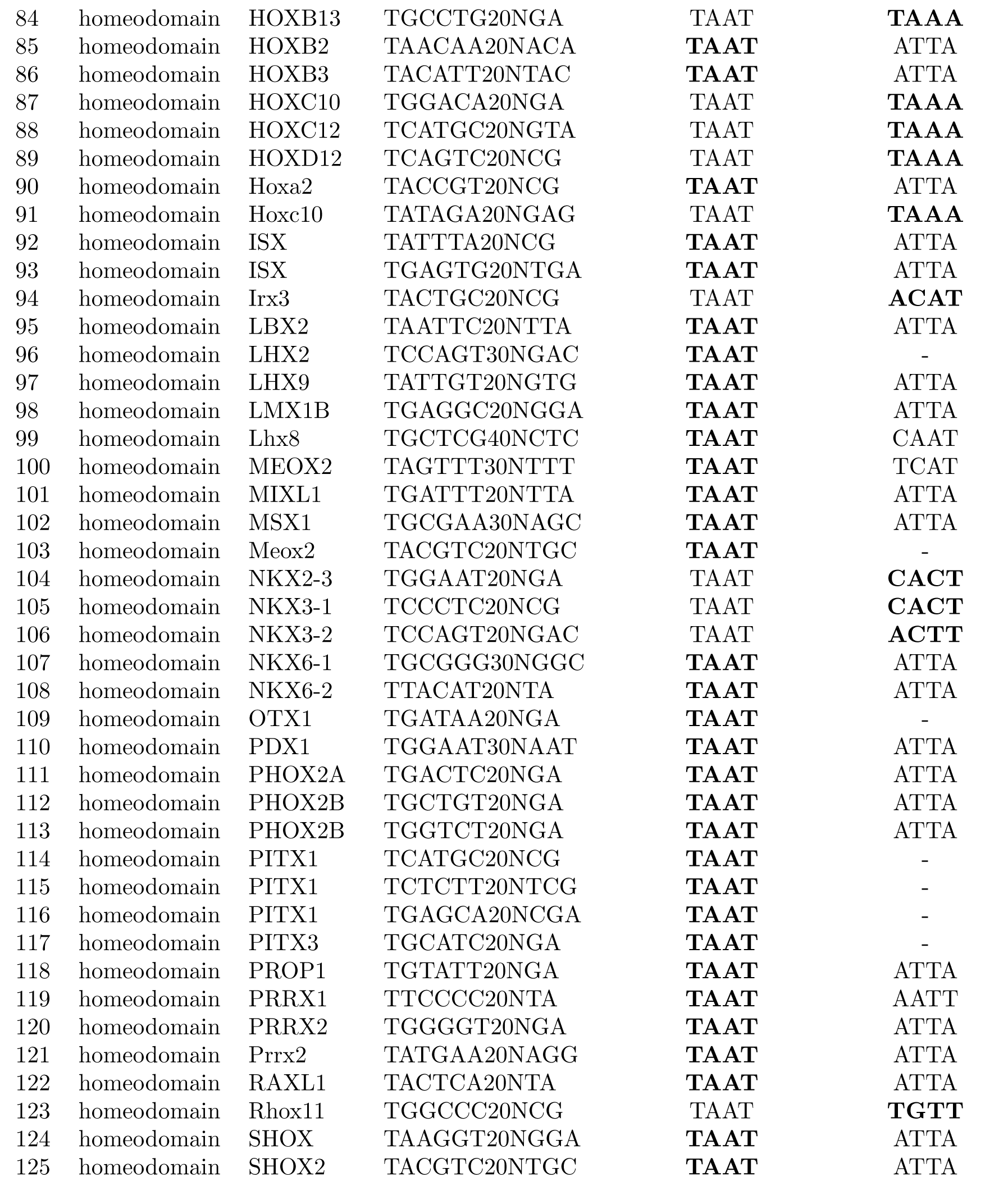
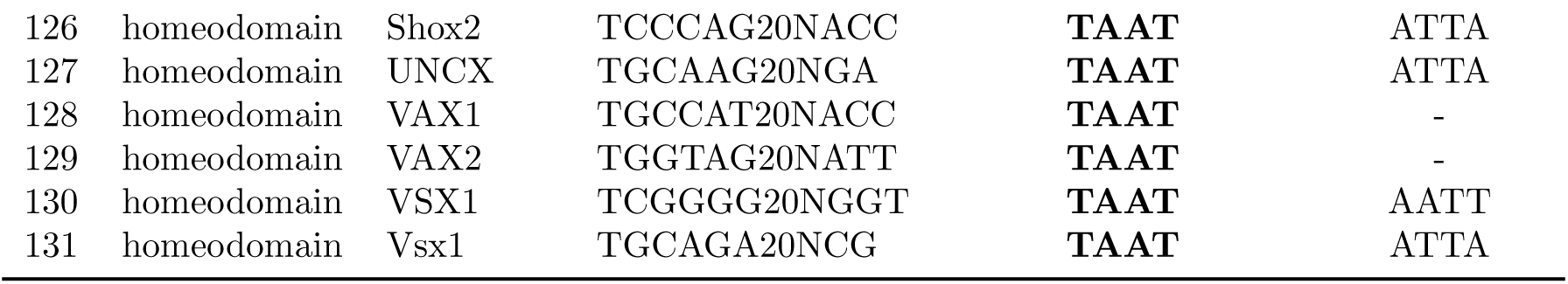
List of successful experiments and the core-motifs used in our analysis for those experiments. Some of the transcription factors in [18] have multiple experiments which are differentiated by the barcode used. If the alternative core-motif is same as the default one we denote it by ‘-’. Between the default and the alternative core-motifs, the one that results in better percentage of aptamers in the competition-test and hence used in our final analysis is shown in bold font.

### S2 Maximum number of mismatches from the core-motif for an oligo to be motif-free

A random oligo of length *n* contains (*n* - *k* + 1) *k*-mers. The probability that a *k*-mer of the random oligo has exactly *d* common bases with the core-motif is 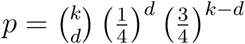 and hence the probability that at least one of the (*n* - *k* + 1) *k*-mers of a random oligo matches with the core-motif at exactly *d* bases is no less than *q* = 1 - (1 - *p*)^*n*-*k*+1^. For the smallest possible *n* = 20 in our database and the smallest *k* = 4 corresponding to the core-motif for homeodomain family, *q* = 0.999909973627, 0.98218056744, 0.557872389117 for *d* = 1, 2, 3, respectively. Thus, setting the threshold *d* = 2, but not *d* = 3 for the maximum number of bases matching with the core-motif ensures that what we call a motif-free oligo is indeed a motif-free oligo. For higher *n* and *k* this threshold remains good as *q* becomes even higher in that case.

### S3 Shape value distributions and discretization cut-offs

One of the important steps of our Co-SELECT algorithm is the discretization of the shape values (see Methods). Instead of using an equal number of discrete levels and equally spaced cut-offs, we first generated the distribution of the values of each of the DNA shape features at all base positions in the oligos from the initial pool and then fitted a mixture of Gaussian distributions for 2,3, and 4 components as shown in Figure S2. We found by visual comparison that the number of components that best fits the distributions are 4,3,4,4 for MGW, HelT, ProT and Roll, respectively. We named the 4 possible discrete levels S,M,H,X for Small, Medium, High and eXtra high, respectively. We determined the cut-offs for the discrete levels by manually inspecting the Gaussian mixture plots and named the cut-offs scheme ‘main-text’ as all our results, unless stated otherwise, uses this cut-off scheme. We also experimented with another variation of the cut-off scheme which we name ‘alternative’, though we did not see drastic change in the results. The exact cut-offs for both schemes are shown in Table S2.

**Table S2:** Discretization levels and cut-offs for the four shape features in two different schemes that we experimented with.

| Scheme name | Shape | Discrete levels | Cut-offs |
| --- | --- | --- | --- |
| main-text | MGW | S,M,H,X | 4.25, 5.4, 5.7 |
|  | ProT | S,M,H,X | 31.9, 34, 36 |
|  | HelT | S,M,H | -8, -5.5 |
|  | Roll | S,M,H,X | -3, 1.2, 5 |
| alternative | MGW | S,M,H,X | 4, 4.6, 5.6 |
|  | HelT | S,M,H | 32, 34.5, 35.5 |
|  | ProT | S,M,H,X | -10.5, -5 |
|  | Roll | S,H | 1.2 |

### S4 Shapemers in motif-free aptamers indeed do not have sequence similarity with core-motif

Our method disambiguates the shape and sequence components of the role of DNA in transcription factor binding by separating the motif-containing oligos from the motif-free oligos. Our analysis establishes the role of DNA shape by showing a significant overlap of shapemers enriched among the two oligo-groups. Thus, our analysis is based on the critical point that the shapemers motif-free oligos no longer carry the the sequence information of the core-motif. To ensure this is indeed the case, we generated the sequence logos of the shapemers in both motif-containing and motif-free oligos from both the initial pool and the pool after round 4. We noticed that the logos among the motif-containing oligos contained the core-motif, as expected, but the logos among the motif-free oligos lacked the core-motif, as postulated. Examples of such logos for the bHLH transcription factor MAX are shown in Figure S3.

### S5 Filter for noisy shapemers

Due to experimental artifacts we found that some aptamers which are neither motif-specific bound nor non-motif-specific bound but are still highly enriched in the final round. The shapemers from these aptamers inadvertently appear to be highly enriched. To avoid such noisy shapemers in each group of motif-containing and motif-free aptamers, we consider in our analysis only those shapemers which originate from at least a fixed fraction of unique aptamers in either group. We call such a fraction a *coverage threshold*. In our experiments we use a coverage threshold of 0.01, i.e., we consider only those shapemers which appear in at least 1 among every 100 unique aptamers in the group.

### S6 Markov model for estimating the probability of the initial pool

One of the critical steps of our Co-SELECT method is the computation of the fold-change or enrichment of shapemers in the final round compared to the initial pool. As shown in Section 2.4, the enrichment of shapemer *x* in the set of oligos *j*, where *j* could be motif-containing or motif-free, is computed as 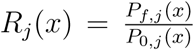, the ratio of the probability of a shapemer appearing in the final pool *P*_*f,j*_(*x*) to the probability of the shapemer appearing in the initial pool *P*_0,*j*_(*x*). Though *P*_*f,j*_ is computed as ratio of the frequency of occurrence of the shapemer in the final pool *H*_*f,j*_(*x*) to the total frequency of all shapemers present in the final pool ∑_*x*_ *H*_*f,j*_(*x*), such a simple method does not work for the initial pool because, as shown in [2], the synthesis of the initial library of double strand DNA sequences is generally biased towards certain oligo sequences or their substrings. Following [2], we use a Markov model that is trained with the shapemers present in the initial pool and then use that model to predict the probability of a shapemer to appear in the initial pool.

For the Markov model there are two options. One option is to train a Markov model for the oligo-sequences and then based on the presence of shapemers in those oligos, derive the probabilities of shapemers for each of the shape features MGW, HelT, ProT and Roll separately from the single Markov model. However, such a derivation of probabilities is complicated and hence we use a second approach which is far simpler. First we generate 4 discretized shape-strings, one for each of MGW, HelT, ProT and Roll, for each of the oligo sequences in the initial pool and then train 4 Markov models separately for the 4 shape features. Furthermore, 4 Markov models are trained for the shape-strings from the motif-containing and motif-free oligos separately. Our shapemers of length 6, and following [2] we use Markov models of order 5.

### S7 Varying enrichment thresholds and discretization levels

We varied the enrichment threshold *ρ* ∈ {1.05, 1.1, 1.2, 1.5} described in Section 2.4 and the two discretization schemes main-text, alternative described in Supplementary Section S3 and noticed that results do not vary for the two discretization schemes. We also noticed that for *ρ* ≥1.5 there not enough enriched shapemers to give us statistical power for drawing any conclusion. On the other hand *ρ* = 1.05, 1.1 is low for a shapemer to be enriched without much noise. Thus we plot the results of the analysis by Co-SELECT in the Figure S4 for all families, both discretization schemes and fixed enrichment threshold *ρ* = 1.2. Each plot in the figure shows two histograms of p-values, one for the experiments (where the shapemers from the motif-containing aptamers for a TF are compared against the shapemers from motif-free aptamers for the same TF) and the other for the controls (where the shapemers from the motif-containing aptamers for a TF belonging to the family under consideration are compared against the shapemers from the motif-free aptamers for a TF in another family) as described in Section 3.2. The area of the bar for pvalue [*x x* + 0.05] for each *x* = 0.00, 0.05, 0.10, …, 0.95 represents the fraction of TFs in the family that resulted a P-value in the range [*x x*+0.05]. The plot also shows *qvalue*_*p*_ for *p* ∈ {0.05, 0.1} that represents the Q-value computed by the R package qvalue corresponding to the P-value *p*. The less the Q-value, the lesser the number false positive if we consider the test to be significant by using the threshold *qvalue*_*p*_.

**Figure S2:**
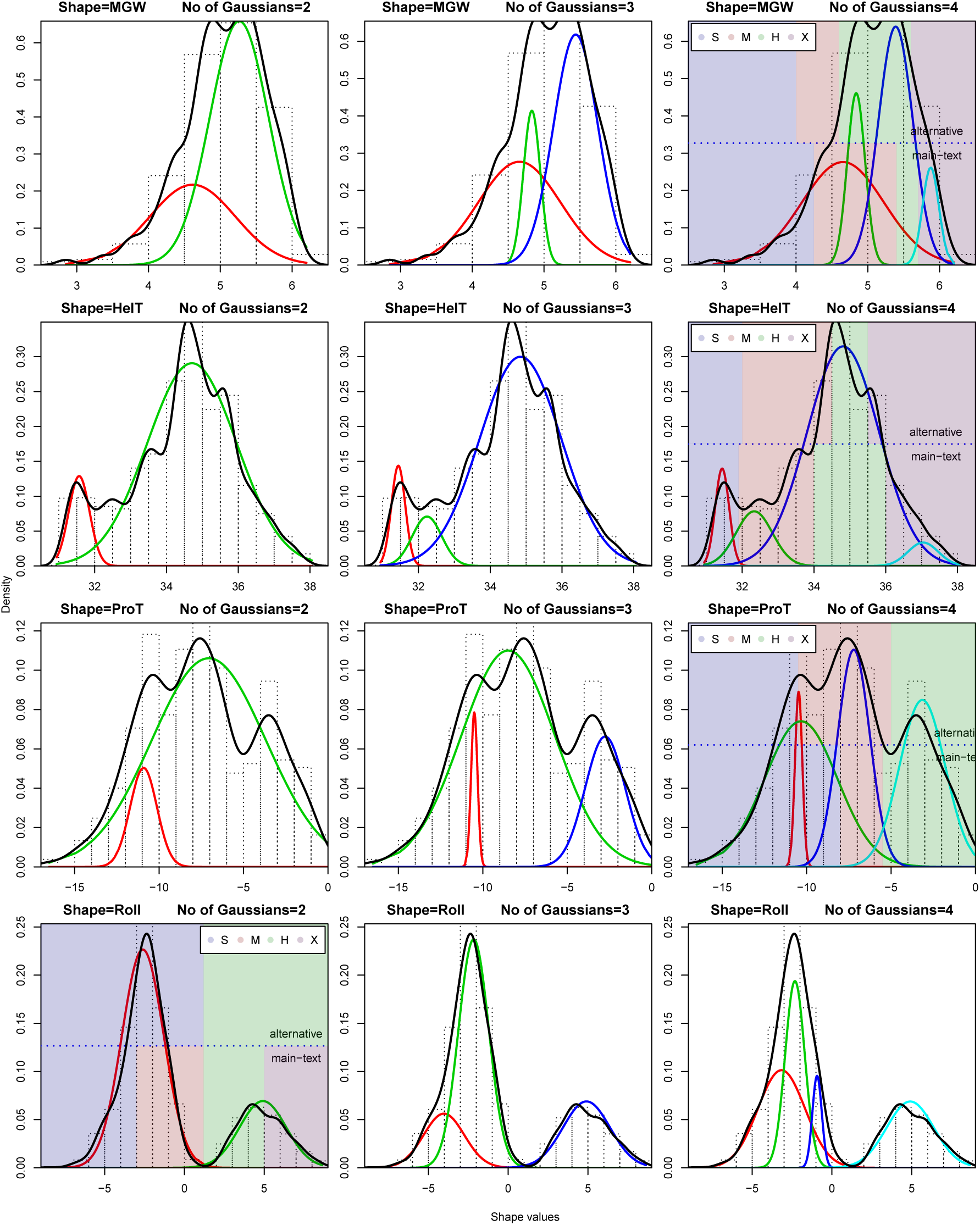
Shape value distributions and discretization levels. Each row corresponds to a shape parameter and each column corresponds to a fixed number of Gaussian components that are assumed to be mixed. In each plot, the black curve represents the density of mixed values and each colored curve represents a Gaussian component as computed using normalMixEM function of R package mixtools [1]. The background of each plot shows the two discretization schemes: bottom shows main-text and top shows the alternative. The different cutoffs and the discretized levels in each scheme are also shown.

**Figure S3:**
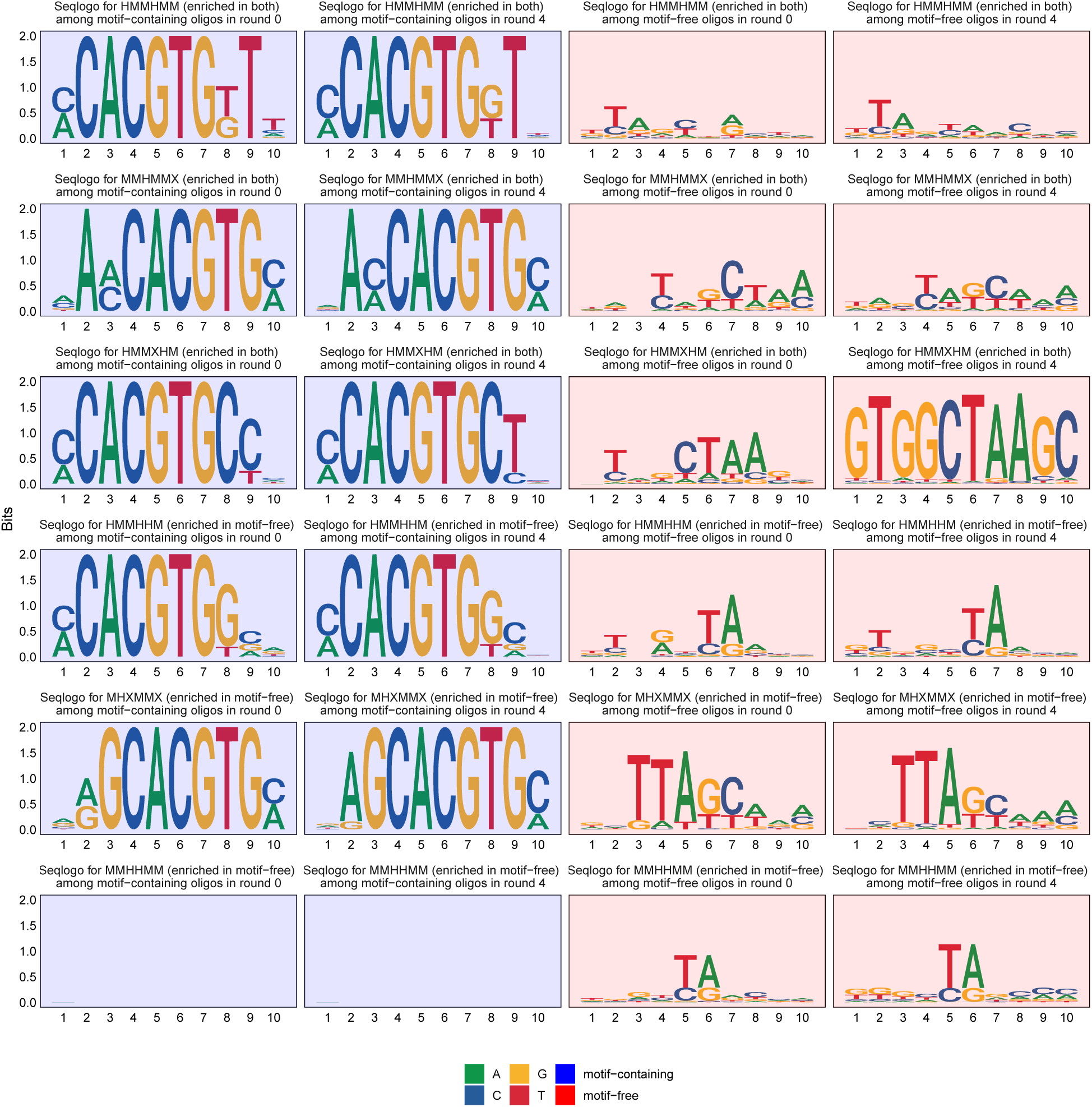
Sequence logo of shapemers enriched in motif-free oligos for bHLH transcription factor MAX. Our analysis revealed that the experiment for MAX resulted in 47 shapemers enriched in the motif-free oligos in round 4. Out of the 47 shapemers, 4 also enriched in the motif-containing oligos in round 4. Here we show the sequence logos of 6 shapemers (3 in each category) among both motif-containing and motif-free oligos in the initial pool and in the pool after round 4. Our shapemers are 6 bases, so we generate the sequence logos of 10 bases considering the 2 flanking bases at both sides as they also contribute in determining the shapemers [20]. For all of the shapemers we notice that the sequence logo does not contain the core-motif CACGTG.

**Figure S4:**
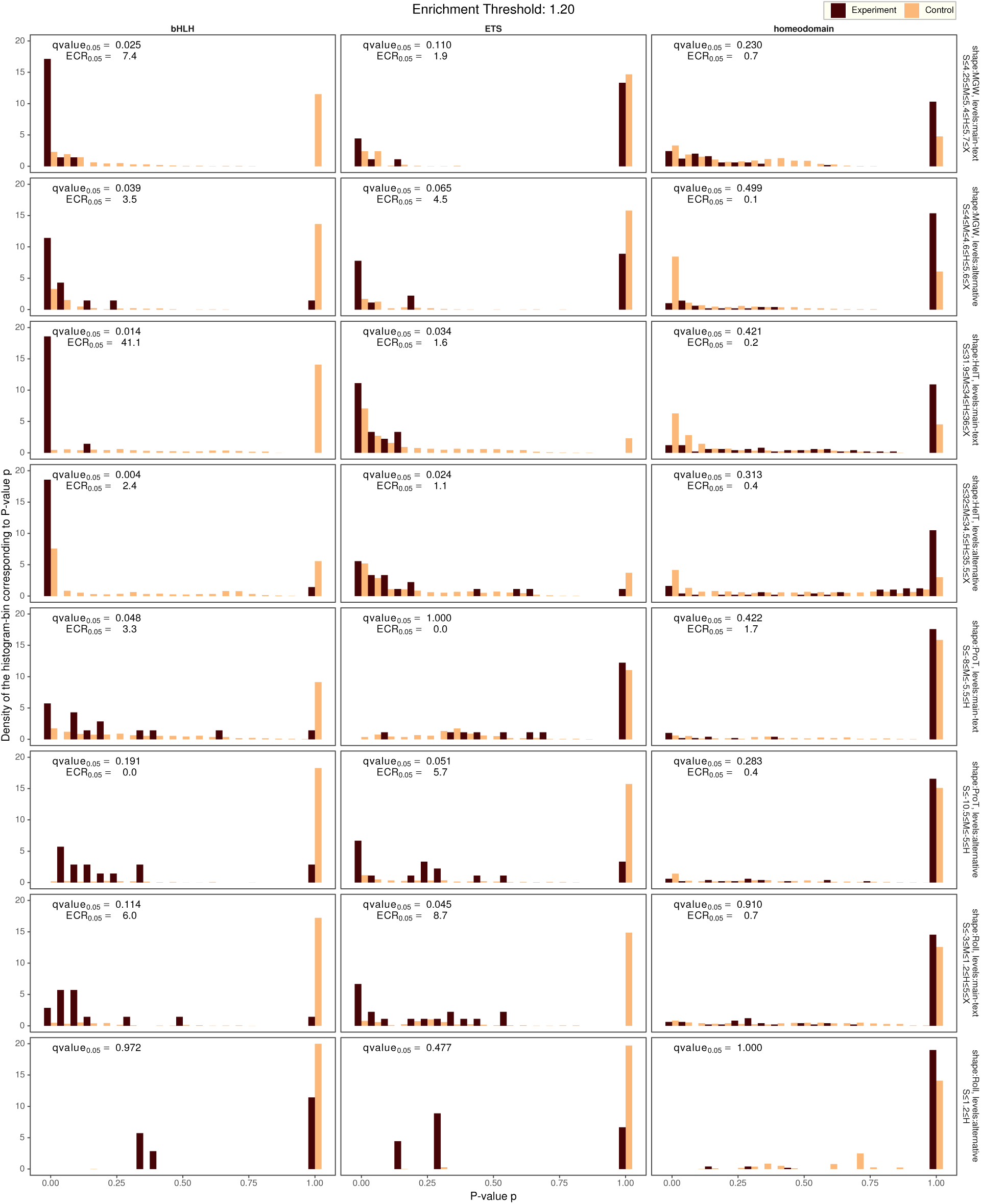
Comparison of normal and control group shapemer for all families and for enrichment threshold 1.20.

